# Human Brain Development: a cross-sectional and longitudinal study integrating multiple neuromorphological features

**DOI:** 10.1101/2022.07.21.501018

**Authors:** Hadis Kalantar-Hormozi, Raihaan Patel, Alyssa Dai, Justine Ziolkowski, Hao-Ming Dong, Avram Holmes, Armin Raznahan, Gabriel A. Devenyi, M. Mallar Chakravarty

## Abstract

Brain maturation studies typically examine relationships linking a single morphometric feature with aspects of cognition, behavior, age, or other demographic characteristics. However, the coordinated spatiotemporal arrangement of morphological features across development and their associations with behavior are unclear. Here, we examine covariation across multiple cortical features (cortical thickness [CT], surface area [SA], local gyrification index [GI], and mean curvature [MC]) using magnetic resonance images from the long-running National Institute of Mental Health developmental cohort (ages 5-25). Neuroanatomical covariance was examined using non-negative matrix factorization (NMF), which decomposes covariance resulting in a parts-based representation. Cross-sectionally, we identified six components of covariation which demonstrate differential contributions of CT, GI, and SA in hetero- vs. unimodal areas. We sought to use this technique longitudinally to examine covariance in rates of change, which highlighted a preserved SA in unimodal areas and changes in CT and GI in heteromodal areas. Using behavioral partial least squares (PLS), we identified a single latent variable (LV; 96 % covariance explained) that recapitulated patterns of reduced CT, GI, and SA that are generally related to older age, with limited contributions of IQ and SES. Longitudinally, PLS revealed three LVs that demonstrated a nuanced developmental pattern that highlighted a higher rate of maturational change in SA and CT in higher IQ and SES females. This novel characterization of brain maturation provides an important understanding of the interdependencies between morphological measures, their coordinated development, and their relationship to biological sex, cognitive ability, and the resources of the local environment.

**Significance:** The complex anatomy of the cortical sheet is best characterized using multiple morphometric characteristics. We expanded on recent developments in matrix factorization to identify spatial patterns of covariance across the cortical sheet. Using a large, well-characterized dataset, we examined the differential contributions of neuroanatomical features to cortical covariation in a single analytical framework using both cross-sectional and longitudinal data. We identified dominant modes of covariance between cortical morphometric features and their coordinated pattern of change, demonstrating sexually differentiated patterns and a strong association with variability in age, socioeconomic status, and cognitive ability. This novel characterization of cortical morphometry provides an important understanding of the interdependencies between neuroanatomical measures in the brain and behavioral development context.

## 1. Introduction

The patterning of the cortical mantle is the result of the development of key features of cortical morphology that are quantifiable using *in vivo* neuroimaging techniques [1,2]. These morphological features, which include cortical thickness (CT), surface area (SA), local gyrification index (GI), and mean curvature (MC), develop at different rates [1] with differing neuroanatomical specificity [3–7]. Thus, studying the relationship between how these morphological properties change through maturation may offer novel insights into normative variation in brain development. There are well-studied morphological changes that occur at specific parts of the maturational time course with known sex differences in spatiotemporal patterning [8]. However, most of our understanding of brain development is derived from studies examining inter-and intra-individual variation in the context of single morphological measures [1,9–23] while ignoring the intrinsic inter-dependencies between measures. Cortical thickness (CT) is linked to the radial (vertical) aspect of cortical growth and is thought to be reflective of synaptic pruning, myelination, and cortical layer cellular density within the radial units and undergoes the most dynamic reorganization in late childhood and through the adolescent period [3,24–27]. By contrast, cortical surface area (SA) peaks between ages 5 and 8 and is associated with the areal expansion of the cortex [3–5] and represents synaptogenesis, rates of neuronal migration, and the density of radial units themselves [3,5,28,29]. Although the underlying mechanisms behind the folding characteristics of the cortex, such as gyrification and curvature, are not as well understood, the imprint of cortical convolutions is observed at the earliest stages of postnatal development [7]. Leading theories for the overall gyrification of the cortex involve physical and biomechanical properties, including cranial pressure and constraint [6], axonal tension [30], and asynchronous expansion of cortical layers [31,32]. At the cellular level, the folding properties are further influenced by neuronal density, the microstructure of the neuronal sheet [33], and the spatiotemporal patterns of neuronal birth and migration [6,31,32,34,35]. Thus, the mesoscopic patterning of the cortical manifold may reflect processes that are cell type-specific, yet potentially interrelated, and specific neurodevelopmental epochs.

Multivariate approaches simultaneously assessing several features may provide further insight to their morphometric inter-relatedness and the true complexity of cortical maturation. Previous studies integrating morphometric measures have estimated subject-specific brain networks derived from combining multiple gray and white matter features [36]. Other studies have used partial least squares correlation to investigate latent variables describing linked patterns of cortical gray matter and white matter covariation [37,38]. Notably, recent studies proposed Morphometric Similarity Networks (MSNs) [2] to elucidate the relationship between multiple dimensions of brain morphology using subject-specific connectomes. They further demonstrated the utility of this approach by relating network-level variation to transcriptomic and cell-specific architecture [39,40]. However, MSNs provide limited interpretability with respect to the specific morphological features that alter connectomic organization (although sensitivity and specificity analysis were included in the original work) [36,39–41]. Further, there have been no clear examinations of longitudinal covariation across measures in these studies.

To address these limitations, we propose a novel implementation of a matrix decomposition technique, non-negative matrix factorization [NMF] [42–46] to study covariance across morphological measures. NMF is conceptually similar to other unsupervised matrix decomposition techniques [47–51] but with a nonnegativity constraint across all inputs and outputs. The recent implementations of this technique in neuroimaging studies [44,45,52–54] have demonstrated the ability to identify spatial patterns of cortical thickness covariance in brain maturation [52]. More recently, this method has been adapted by our group to enable the integration of multiple structural metrics in the context of the human hippocampus [46], striatum [55], and cortex [56] as a means to examine the relationship between inter-individual neuroanatomical covariance and variation in demographic and behavioral features. Here, we use NMF to study cortical neurodevelopmental covariance, cross-sectionally and longitudinally, across CT, SA, GI, and MC. We further related these patterns to age and sex, socioeconomic background, and cognitive ability to better understand inter-individual differences in cortical patterning and maturation. Finally, to assess the putative functional relevance of our components, we assessed how the identified morphometric covariance patterns situate along the maturational stages of gradients of functional connectivity within the different periods of childhood, adolescence, and adulthood [57,58].

## 2. Results

### 2.1. Workflow Overview

We included a subset of the large-scale longitudinal T1-weighted magnetic resonance imaging (sMRI) data from the National Institute of Mental Health cohort (NIMH; Bethesda, MD, USA), a longitudinal study of brain development. The final sample consisted of 776 participants and 1142 scans in total from participants aged 5 to 25 years old (see section 4.1): 776 participants for cross-sectional analysis (357 F; mean age:12.4, Standard Deviation [SD]:3.49) and 183 participants for longitudinal analysis (77 F; three repeated scans per subject, approximate interval 2.8 years; mean age: 11.2, SD: 2.7) (see Table.S1 and Fig.S1 for both cross-sectional and longitudinal samples characterization) after stringent quality control procedures [59]. All morphological features were derived from sMRIs preprocessed with the CIVET pipeline (version 2.1.0) [60,61] (Fig.1A) to examine cross-sectional and longitudinal patterns of morphometric covariance using NMF. In the cross-sectional analysis, we used each subject’s cortical measures using their first scan. In the longitudinal analysis, we extracted age-related individual slopes as a proxy of change in the cortical measures over time to investigate the pattern of *coordinated maturation* of cortical features. Further, we examined how these covariance patterns relate to age, sex, intelligence quotient (IQ; estimated using Wechsler Abbreviated Scale of Intelligence (WASI)), and socioeconomic status (SES; quantified using the Amherst modification of the Hollingshead two-factor index [87,88]; see supplementary section S1), using univariate measures to capture group-level trends, and multivariate methods (PLS) to examine the relationship across demographic variables (Fig.1C). Finally, we assessed the position of identified components along the unimodal-transmodal axis of stages of cortical functional connectivity gradient maturation (childhood, adolescence, and adulthood; see methods section 4.6) (Fig.1D) to better understand their putative functional relevance.

**Fig.1.**
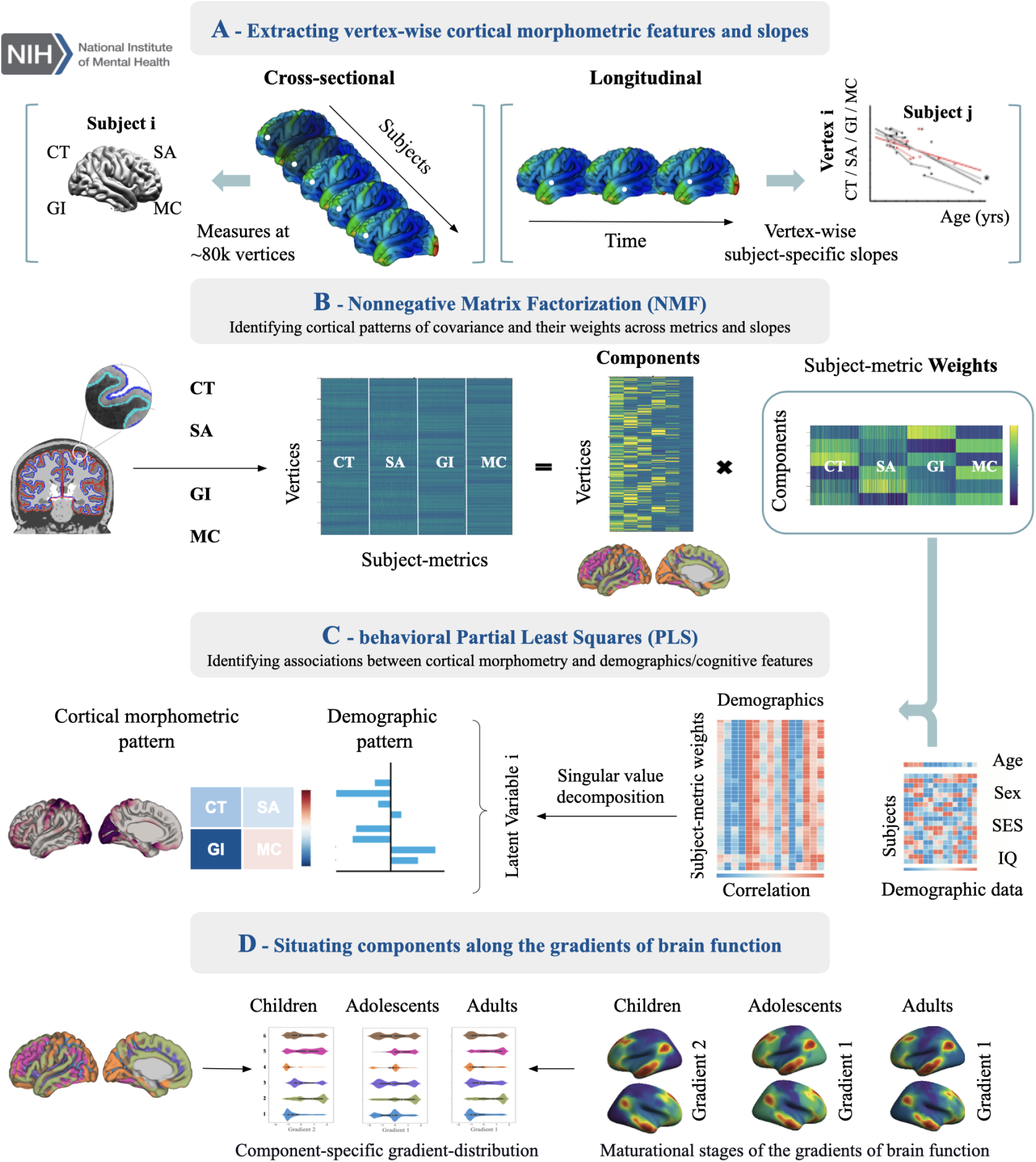
Workflow. **A)** Four cortical metrics (CT, SA, GI, MC) were extracted, and cortical vertices of all subjects were concatenated in columns to build an input morphometry matrix. For longitudinal analysis, age-related slopes were extracted using linear mixed-effects modeling. **B)** NMF decomposes the input matrix into a components matrix, representing spatially distinct components of covariance of morphometry across subjects, and a weights matrix, representing the extent to which each subject loads onto the identified components. The optimal number of components was selected by balancing the accuracy and spatial stability of decomposition by performing a stability analysis (see section 4.4). **C)** Behavioural PLS (bPLS) was performed to identify patterns of covariance across components and their specific morphometric weights with demographic data (Age, Sex, SES) and cognitive ability (as indexed by IQ). **D)** we assessed the position of identified components along the stages of the functional gradients’ gradual maturation (childhood, adolescence, and adulthood) as described by Dong et al. [57].

### 2.3. Morphometric Covariance Results

#### 2.3.1. Cross-sectional morphometric covariance: Cortical patterning

Orthonormal projective nonnegative matrix factorization (OPNMF) provides a set of parts-based and spatially orthogonal patterns of neuroanatomical covariance across vertices (which we refer to as components throughout the remainder of this paper; Fig.2A), and describes the inter-individual variation in these patterns using subject-level weights for each of the morphometric measures (Fig.2B). The number of components is chosen by the user. Here we examined stability and reconstruction error [46,55] to inform our decision on the number of components chosen (See methods 4.4 and Fig.S2 in supplementary).

**Figure 2).**
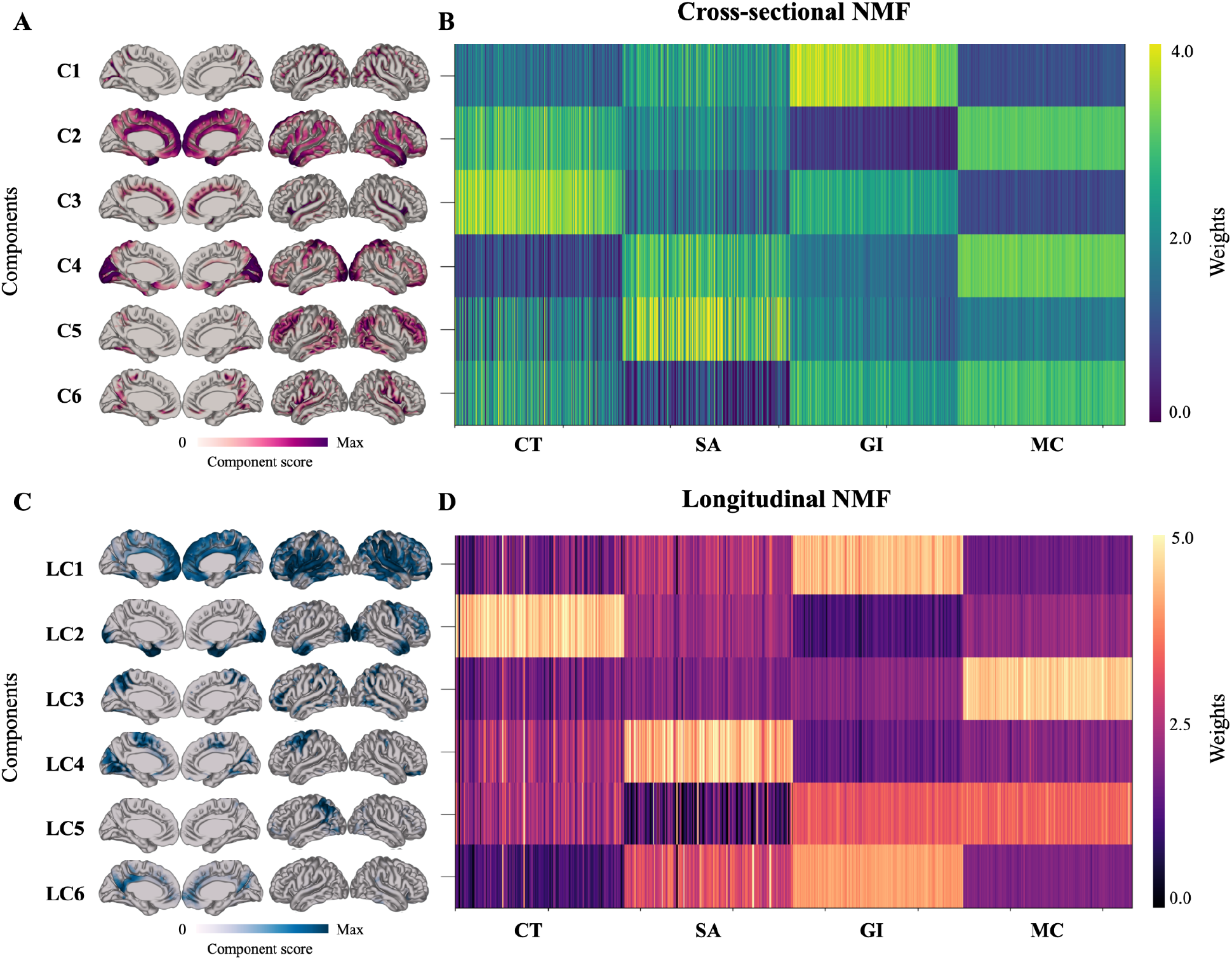
**A & C**. Spatial cortical components of a 6 component decomposition solution. Identified components are projected onto an average template brain. Lateral and medial views of both hemispheres are shown. Darker blues correspond to higher component scores. **A)** Vertices that are grouped together in components share the same pattern of morphometry, while in **C)** vertices that cluster together share a coordinated pattern of maturation across four metrics. **B & D.** The weight matrix shows a comparative morphometric pattern of each component (z-scored across rows and shifted by minimum value for better visualization). In **B),** brighter colors represent higher values (measurements), while in **D)** brighter colors represent higher slopes and preservation. Note that the weight matrix is normalized by z-scoring within each row for the purpose of visualization to better capture the comparative morphometric features of each component.

Our analysis of the NMF outputs identified six stable morphometric components (Fig.2A) in the cross-sectional dataset (see Methods 4.4 and supplementary Fig. S2). Notably, even though there were no topological constraints in our analysis, there is significant topological specificity and bilateral representation for each component:

1. Component [C]1 is characterized by higher weight values of GI and SA, primarily in the sulcal depth of temporal, occipital, and parietal association cortices.
2. C2 is characterized by higher values of CT and MC, moderate SA, and lower values of GI in limbic and heteromodal regions along the medial surface, extending along the temporoparietal junction, Broca’s, and Wernicke’s area to the orbitofrontal cortex, and the superior gyrus.
3. C3 is characterized by high CT, moderate SA, and GI weighting across the insular cortices and cingulate sulcus.
4. C4 is characterized by higher MC and SA weight values along the fusiform and orbitofrontal gyri and unimodal regions along the lateral and medial occipital lobe and postcentral gyri.
5. C5 is characterized by primarily higher SA weights in primarily heteromodal regions of the inferior temporal, inferior parietal, and dorsolateral medial prefrontal cortex.
6. C6 is characterized by low SA, moderate CT and GI, and high MC. This component includes cortical regions of pre and postcentral gyri (primary sensory and motor cortex) and precuneus.

#### 2.3.2. Longitudinal morphometric covariance: Coordinated change

Longitudinal analysis using NMF also identified six stable spatial cortical components (see Supplementary Section S8 for constructing the input matrix and Supplementary Fig.S2 for stability and reconstruction error analysis) representing a selection of vertices sharing a coordinated morphological pattern of *change* across the four metrics (Fig.2C). Unlike the cross-sectional analysis, here the weights matrix describes the extent to which each subject-metric change (i.e., age-related slope) loads onto the identified spatial pattern (i.e., components) (Fig.2D). Higher values (i.e., less negative slopes, or positive slopes) indicate preservation, while lower values (i.e., more negative slopes) indicate the higher rate of decrease of a metric. Fig.2B shows the subject weight matrix, describing morphometric patterns associated with each component. Like the cross-sectional analysis, we observe significant bilateral symmetry, but this analysis yields a different topological result relative to the spatial topology of the cross-sectional results:

1. Longitudinal Component [LC]1 describes relative preservation of GI, dominant decline of CT, and a moderate decline of MC and SA across much of the cortex, in keeping with the known literature on brain morphology changes during development [1].
2. LC2 describes relative preservation of CT, a sharp decline of GI, and a decline of SA and MC in bilateral occipital and temporal poles and the right precentral gyrus/ primary somatosensory areas.
3. LC3 describes the preservation of MC and a decline of CT, SA, and GI in the sulcal depths and precuneus.
4. LC4 describes relative preservation of SA and a steeper decline in GI and moderate CT and MC decline in unimodal areas such as primary somatosensory areas and in the cuneus and lingual gyrus.
5. LC5 describes a steep SA decline and moderate CT decline in the left temporoparietal junction.
6. LC6 describes a steep CT decline and a moderate MC decline in the posterior cingulate gyrus.

### 2.4. Post-NMF analyses results

We further investigated relationships across the patterns identified by our morphometrics and demographics using multiple linear regression models and behavioral partial least squares analysis (bPLS) [46,62]. bPLS identifies patterns of variables that maximally covary across two sets of data through latent variables. This method has been conventionally used to relate a set of neuroimaging data to a set of behavioral data [46,62–64]. Here, we modified the method to examine the patterns of associations across 1) subject-level NMF weights of cortical morphometric measures and 2) demographic data through latent variables (LV). Each LV consists of three parts: a brain score and a demographic score, describing the contribution of each of the variables to the LV, and a singular value, indicating the percentage of total covariance that is described by the LV [46,62–65] (see Methods section 4.5 and supplementary section S10.2).

#### 2.4.1. Multiple linear models regression results (Cross-sectional NMF)

To investigate the specific sex and age-by-sex associations with regional patterns of morphometric variation, we also performed multiple linear regression analyses (see Table. S2 for statistical results, Fig. S3 for components’ weights plotted against age for males and females, and S10 for methodological details). All component weights were significantly (corrected for multiple comparisons across components; significance threshold: p<0.008) decreasing with age, except for MC weights showing significant increases in C1, decreases in C2, and no association with age in C3-C6. Subject-specific weights across many components and metrics were significantly greater in males, such that SA weights across all components, CT across components 4 and 6, and GI across components 2, 5, and 6 were greater in males than females. Conversely, females’ weights for MC were significantly higher across all components. Significant age and sex interactions were observed for SA weightings in components 1, 4, 5, and 6 such that males showed a slower SA loss over development relative to females. Significant age and sex interactions were observed for CT weights in component 4 such that males showed a slower CT thinning relative to females. Socioeconomic privilege revealed positive associations exclusively with GI weights across components 1, 2, and 5. IQ scores, however, were not associated with cortical weights.

#### 2.4.2. Cross-sectional NMF analysis results are mostly associated with age

PLS identified a single significant LV (p<0.05), explaining 94.6% of the covariance across cortical and demographic data (Fig.3). The largest contribution to the demographic covariance patterns is clearly age. In the morphometry pattern, we observe a dominant contribution of GI reduction with age across all components, as well as CT thinning, SA reduction, and a local increase in the sulcal depth in MC. However, we also observe the subtle impact of lower SES and IQ scores on these patterns. These findings are generally in line with previously described age-related changes, particularly the linear trajectory of pronounced GI reduction in widespread cortical regions of precentral, temporal and frontal areas [66].

**Figure.3).**
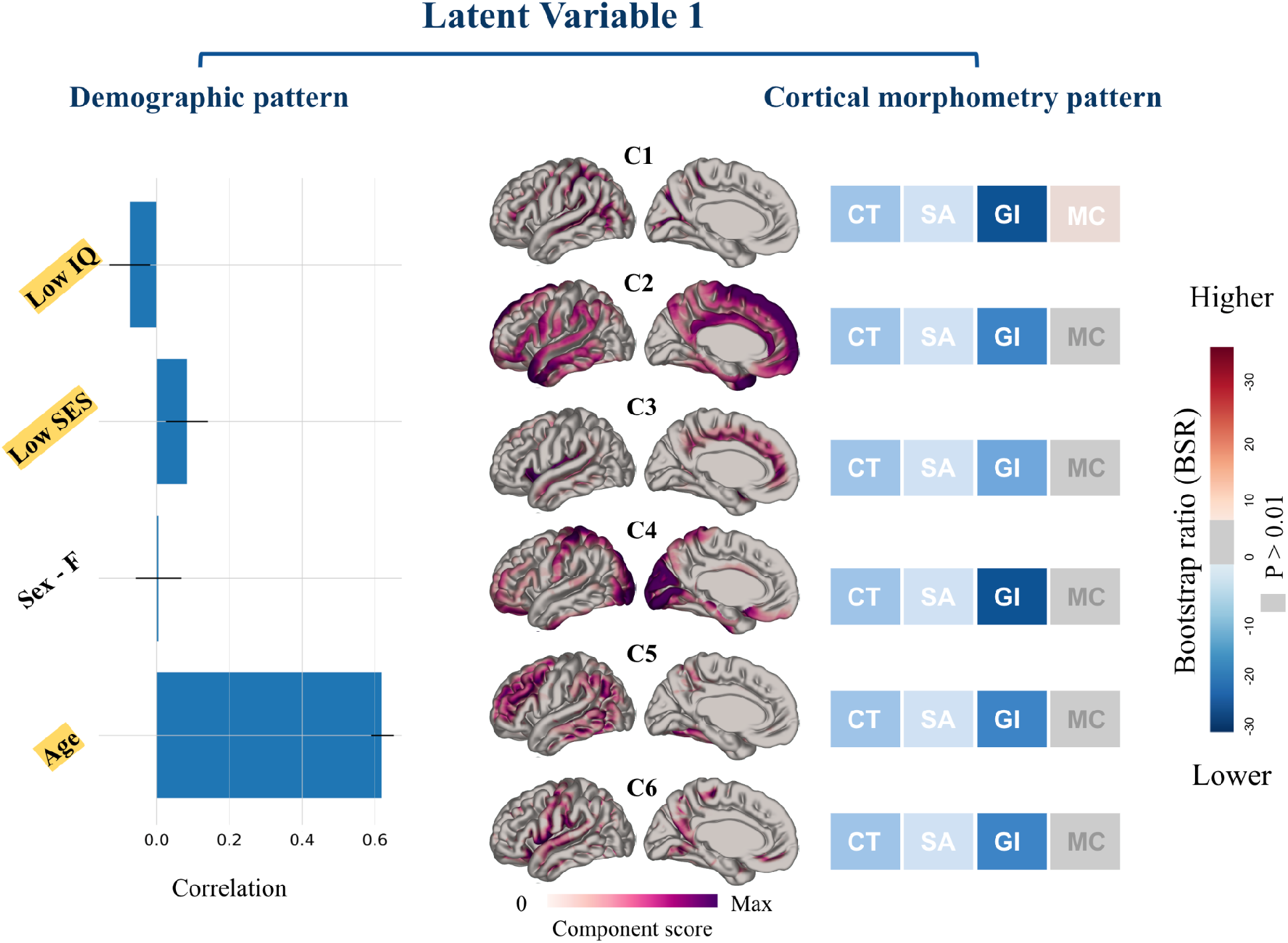
The single significant LV describing cross-sectional NMF and individuals’ characteristics. Cortical components of morphometric features (right) and demographic data (left) contributing to the LV are shown. **Right)** The brain maps are summarized with their morphometric profiles, each with a set of four metrics corresponding to a single spatial component; for each significant morphometric feature contributing to the LV, the bootstrap ratios are displayed on the components’ morphometry profile, in which lower values of a metric is color-coded in blue whereas higher values, in red. **Left)** The bar plot describes the contribution of demographic variables to the identified LV. The x-axis demonstrates the correlation of each demographic variable in the LV. Error bars indicate the 95% confidence interval; variables with a BSR> 1.96 (p <0.05) are described as contributing to the LV [46] (color-coded in yellow). Please note the opposite directionality of the Hollingshead score used in the analysis and figures with the SES.

#### 2.4.3. Multiple linear models regression results (Longitudinal NMF)

Multiple linear regression analyses were also performed to determine the statistical significance of the regression models using age-related slopes and individual characteristics (see Supplementary: Table.S3 for statistical results). Briefly, NMF-derived GI slope weightings were significantly (corrected for multiple comparisons across components; significance threshold: p<0.008) positively correlated with age in components LC2 and LC4. In contrast, SA weights were negatively correlated with age. As GI, CT, and SA generally decrease with age, in the context of the longitudinal implementation of slopes, a positive association between GI slope and age translates into a lower rate of change (i.e., the slope is moving towards zero). These findings, therefore, suggest that cortical complexity (as indexed by GI) decreases at a more stable rate with age (i.e., the older the age, the lower the rate of GI changes with time) while the decrease in other morphological measures (SA) is more pronounced with age. SA in components LC3 and LC4 and MC in LC2 showed sex-specific correlation patterns. Significant age-by-sex interactions were observed across SA, CT, and MC weightings exclusively in LC2 in bilateral occipital and temporal poles and the right primary somatosensory areas. Notably, only SA weights in LC4 were significantly correlated with SES.

#### 2.4.4. Longitudinal NMF analysis results demonstrate increased specificity to demographics

bPLS identified three significant (p<0.05) LVs that together explained 97.6% of the covariance across cortical and demographic data. In Fig.4, cortical morphometry and demographic patterns contributing to the LVs and their corresponding bootstrap ratios (BSR) are displayed. LV1 (p = 0.0001), explaining 47% of covariance across the brain and demographic data, describes a sexually differentiated pattern of change in the cortical morphometry slopes of five of the six cortical components (LC1 - 4 and LC6). These results, in line with the linear regression results, suggest that more socioeconomically advantaged and high IQ females accelerate through the known neurodevelopmental processes. These maturational changes mostly involve SA (LC1, 3, 4, and 6) and CT (LC1 and 2) reductions in specific components, almost uniquely in regions that are related to heteromodal processes. MC changes in LC6 exclusively, show a more stable or increasing rate of change in females and lower SES demographic groups as they mature (Fig.4A).

**Figure.4).**
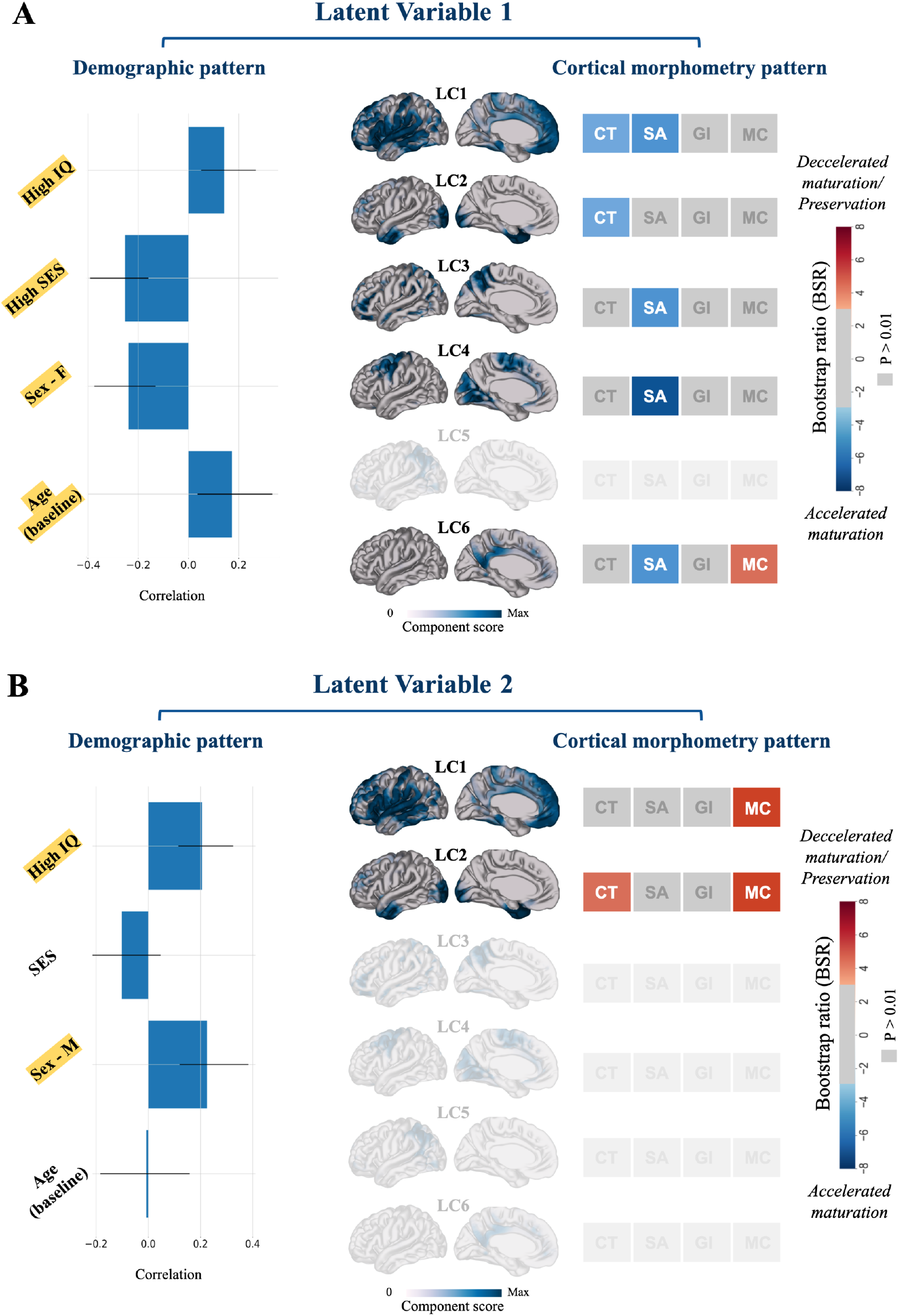
Latent variables identified by bPLS. bPLS analysis identified three (LV1: A; LV2: B; LV3: supplementary Fig.S4) significant latent variables (p<0.05), each identifying a pattern of linear correlation between NMF weights and demographics. Bar plots on the left describe the contribution of demographic measures in which the x-axis demonstrates the correlation of each demographic variable within an LV. Error bars indicate the 95% confidence interval; only variables with a BSR > 1.96 (p <0.05) are described as contributing to each LV [46] (color-coded in yellow). For each bar plot, cortical components contributing to the LV are shown. The morphometric profile of each component describes to what extent changes across a given metric are identified as being accelerated (blue) or decelerated (red) in the spatial component in relation to the demographic pattern shown in bar plots. Only cortical variables with a BSR > 2.58 (p <0.01) are described as contributing to a LV (color-coded according to the BSR plot shown on the right).

LV2 (p = 0.005), explaining 26% of covariance, describes a pattern that is more component-specific, loading predominantly onto LC1 and LC2. This LV reveals a pattern in which male sex and higher IQ are associated with changes in MC and CT in two of the cortical components (LC1, LC2). The relationship is such that male individuals with higher IQ show relative preservation of MC (in LC1 and 2) and CT (in LC4) compared to females and lower IQ participants as they mature through adulthood (Fig.4B).

LV3 (p = 0.02), explaining 24% of covariance across the data (p-value = 0.005) describes a specifically age-related pattern of GI change in two cortical components, LC2 and LC4, such that GI slopes show a milder decline in older ages (i.e., the slopes of GI change become less negative in older ages; see Fig.S4). Together, these LVs suggest that patterns of coordinated anatomical change are sexually differentiated, influenced by environmental factors such as SES, and related to different cognitive abilities. Contrasting this with the cross-sectional analysis allows us to infer how coordinated measures of brain change, a proxy for brain development, influence sex and environment, suggesting that maturational processes are sensitive to these factors.

### 2.5. Morphometric components occupy different positions along gradients of brain function

Previous studies have shown that morphological networks can be contextualized by comparing their localization relative to functional networks along the unimodal-transmodal gradients [36,67–69]. In our study, we sought to investigate how the morphological components that we observed align with recent reports of age-related reconfiguration of functional gradient organization developed by Dong et al. [57]. In children (<12 years old), the functional gradients are “anchored” in the unimodal cortices (somatomotor and visual cortices), which, in adolescence (>12 years), transition towards an adult-like spatial organization, with the default network-related regions emerging at the opposite end of the gradient space from primary sensory and motor regions [57]. We, therefore, assessed the correspondence of the identified morphometric covariance patterns with gradients of connectivity for adolescents and adults and the second gradient for children to capture the same cortical hierarchy across all three age groups. Fig.5 demonstrates parcellation brain maps for NMF components, demonstrating the gradient value distributions of vertices within each component for the three age groups in which positive values indicate proximity to the transmodal end of the gradients.

**Figure.5).**
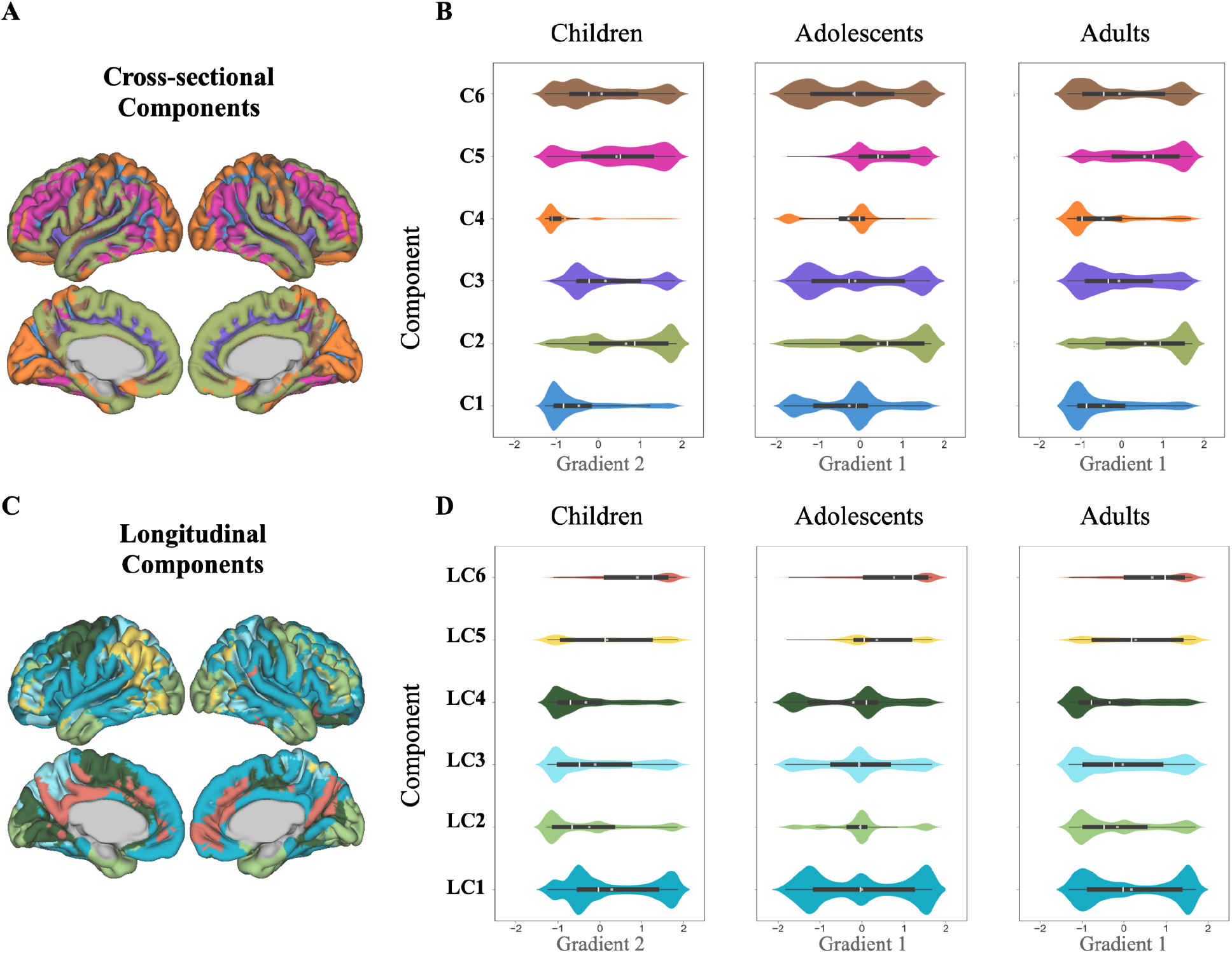
**A&C.** Distribution of cortical components derived from NMF for cross-sectional (A) and longitudinal (C) analyses. Gradient value distributions of vertices within each component’s spatial boundaries were extracted from the maps of the second gradient of functional connectivity for children (6-12 years old), and the map of the first gradient map from adolescents (12-18 years old; obtained from Dong et al. [57]) and adults (22-35 years old; obtained from Margulies et al. [58]). **B&D.** Violin plots demonstrating z-scored gradient value distributions from the three age group maps for each NMF-derived component, where more positive values indicate proximity to the association end of the gradient. The white line in the box plots indicates the mean, and the gray box indicates the median. The Connectome workbench [70] was used to generate the NMF parcellation brain maps (A&C).

As in [57], we used the second gradient in the < 12-year-old group to represent the transition from unimodal to heteromodal described in the literature [58]. The cross-sectional NMF components almost occupied one of two ends of the gradients relative to the three age groups. C2 and C5 are associated with a position along the association end of the gradients defined for each age group, while components C1, C3, and C4, which are more sensorimotor, lie towards the sensorimotor gradient end, and C6 is prominently distributed over the two ends of gradients. Comparing the distribution of components along the second gradient of children relative to the adolescent first gradient reveals a pattern of shifting mean and median values (Fig.5) towards the transmodal end of the gradients as seen in C1, C4, and subtly in C6 (which are generally unimodal-centered components). This is followed by the segregation of these components once more during adulthood.

In the longitudinal NMF (coordinated maturation), components’ position demonstrated a slightly different pattern than the cross-sectional analysis. LC1, which is situated in posterior and anterior cingulate regions, demonstrated a mostly stable heteromodal configuration relative to the three age groups. Similarly, LC2, which represents parts of the visual stream, was also quite stable. LC2, LC3, and LC4 occupied a more unimodal position in relation to the children’s map, with a shift in distribution during adolescence and a return to a segregated representation during adulthood. Our findings suggest that our components may have functional relevance to areas undergoing refinement and that are reconfiguring to enable the integration of unimodal and heteromodal regions.

## 3. Discussion

The present work uses a multivariate framework to examine morphological and maturational covariance patterns across measures of CT, SA, GI, and MC. The group-level covariation captured in our analysis speaks to inter-individual variation, is mostly attributable to coordinated maturation across morphological measures, and is impacted by sex and social advantage. Unlike the longitudinal analyses, cross-sectional variation in neuroanatomical patterns is mostly attributable to age. Finally, we observed that cortical components occupy different positions along the sensorimotor-association axis of brain function maturational gradients.

### Identifying regions of cortical variability and the choice of parcellation

In our PLS with cross-sectional NMF components, we observed a dominant pattern that generally captured known age-related associations. However, our PLS analysis with longitudinally-derived NMF components demonstrates that cortical maturation rates have morphometric specificity that is related to age, sex, cognitive ability, and social advantage. This latter finding is in line with the previously described sexually differentiated patterns of cortical maturation (i.e., cortical thinning, surface area reduction) with females maturing earlier [71,72] and demonstrating significantly higher rates of cortical change through maturation [73,74] in heteromodal regions such as the temporal, temporoparietal, and orbitofrontal cortices. The differing tempo of maturation has also been previously reported in Raznahan et al. [1], such that sex differences captured across the global cortical volume were mainly reported to be driven by sex differences in SA and, to a lesser extent, CT [1] while GI has been reported to show only subtle sex differences with age in localized frontal regions [74], suggesting that the mechanisms underlying age-related changes are distinct. These sex-differentiated patterns are thought to be genetically determined [73] and could be explained by the different timing and rate of fundamental biological maturation moderated by hormonal processes [75,76].

Our inclusion of socioeconomic advantage and cognitive measures provides some insight into how the local environment may impact brain maturation. These findings complement recent work in this area. A recent PLS analysis investigating the association between the whole-brain structural connectomic organization and childhood SES [77] demonstrates that the structural connectome mediates the relationship between SES and cognitive ability. In line with this, our PLS analyses revealed a pattern of lower SES and IQ associated with thinner CT, reduced SA, and reduced complexity in our cross-sectional PLS analysis.Higher SES and IQ were also associated with accelerated cortical thinning and area reduction, which can cautiously be interpreted as an acceleration in maturational trajectories and is consistent with the previous literature [78,79].

Seidlitz et al. [2] previously demonstrated that inter-individual variation in topography of multiple dimensions of brain organization in the MSN framework is predictive of the inter-individual variation in IQ [2]. PLS analysis between the MSNs’ nodal degree and IQ measurements revealed patterns of a positive association between these measures in the left frontal and temporal cortex (left-lateralized temporal and bilateral frontal cortical areas) with higher general IQ. They also demonstrate that a higher degree in bilateral primary sensory cortical areas was specifically predictive of a higher nonverbal IQ [2]. These results are in alignment with the pattern of cortical morphometry-cognitive and demographic associations captured through our bPLS using longitudinal variables. LV1 was associated with higher IQ and SES, and female sex has been linked to covariation of accelerated thinning in cortical thickness (in LC1) and surface area reduction in LC1 and LC4.

Taken together, the results highlight the complex dynamic changes occurring across maturation and emphasize that inter-individual differences significantly influence the normal variation in cortical patterning. Comparing the results from the two PLS analysis of cross-sectional and longitudinal NMF weights emphasize that inter-individual variability in factors such as age, sex, environment, and cognitive ability are better explained by variations in the tempo of anatomical change than cross-sectional morphometric covariation [1]. Overall, these findings support that the integrative framework we have proposed simultaneously captures information about multiple dimensions of brain morphometry that meaningfully explain a significant proportion of inter-individual variance in age, sex, IQ, and SES.

### Spatial alignment of NMF components with the maturational stages of functional gradients

Human brain development follows a pattern of an early maturation of the unimodal visual and sensorimotor areas, followed by refinement of the multimodal association cortices [10,19,57]. Leveraging the previously described age-dependent patterns of gradual maturation in the macroscale of cortical organization, we investigated how NMF components situate along the gradients of brain function across development. From the point of view of the transition in the functional hierarchy, the second gradient in children demonstrates a shift from a more unimodal-centered organization to a transmodal anchored framework. However, such developmental transitions are not present in all components, such as (C2, C3, C5) and (LC1, LC5, LC6). Their positions were spatially consistent across children (second gradient) and adolescents (first gradient);, this may highlight the presence of stable features of the cortical macroscale organization of the cortex in this overarching pattern of developmental transition [57]. From adolescence to adulthood, components show a pattern of a shift from an overarching mid-centered distribution in the first gradient (visual system) to a more multimodal-centered organization towards the extreme ends of the gradients. These findings could potentially reflect the continued refinement of the visual system within the global connectivity structure through young adulthood [57]. Taken together, these age-dependent and interconnected patterns of shift across the position of morphometric-derived components along functional gradients may reflect the refinements towards the facilitation of multimodal information integration and “segregation of local, specialized processing streams” in the adult-like cortical architecture [57].

### Limitations

The findings of this study should be interpreted while considering its limitations. The generalization of our results is limited as NMF identifies data-driven components; subsequently, the identified spatial patterns are specific to this dataset, and the age range studied. Another limitation is the linearity presumption in multivariate approaches. PLS, for example, reduces the complexity of analyses by providing a concise summary of the data and may mask potential non-linear relationships. The same is true for the use of slopes as the proxy of change in our longitudinal analysis; our analysis of coordinated maturation is restricted to modeling the effect of age on cortical changes in a linear fashion, which conceals nonlinear changes. However, some neuroanatomical changes are thought to occur non-monotonically [10,80,81]. For example, SA peaks at the age of 8 and decreases gradually afterward; through the linear measure of slope, this cubic trajectory could be interpreted as a slow decline with age instead. A caveat of the current work could therefore be the large age range studied that might conceal interaction effects that vary by age, such as the cognitive- and demographics-related variability (as shown in previous studies [82]). While NMF is an ideal method to explore large population samples by providing group-level covariation networks, it is limited by the inability to construct individual-level networks; this should be borne in mind for the future clinical implications of this framework. Lastly, manual quality control could potentially be subject to errors and inconsistencies.

### Future directions

This work would benefit from the integration of white matter indices (such as T1/T2, T2*, FA, and Mean diffusivity (MD)) from multi-modal data to delineate networks of coordinate maturation across gray and white matter boundaries, enabling better characterization of the complex and dynamic codependencies between gray and matter tissues. Additional cortical metrics to be investigated using higher resolution MR images could be boundary sharpness coefficient, which has been proposed as a proxy to capture alterations in microstructure at the cortical gray/white matter boundary [83,84], and measures obtained from different cortical depths, that have been shown specificity in capturing alterations in neuropathological conditions [85–88]. It would also be important to ascertain whether the morphometric components of covariation map onto the spatial expression of certain cell types, gene expression (transcriptional profiling of spatial components), evolutionary hierarchy, and developmental expansion [89] profiles. A more nuanced characterization of this type would potentially provide a mechanistic understanding of morphological development and lead the way towards understanding neurodevelopmental disorders.

### Conclusion

Leveraging a data-driven technique, we identified patterns of cortical covariation across integrated measures of cortical morphometry and their synchronized rate of maturation in typically developing youth. Taken together, we observed a non-uniform relationship between morphometric measures throughout the cortex underlying fundamental neurodevelopmental processes that covary together. The identified patterns were age-related, sexually differentiated, influenced by individual differences in socioeconomic factors, and associated with cognitive ability. This novel characterization of cortical morphometric maturation provides an important understanding of the interdependencies between morphological measures, their coordinated development, and their relationship to critical factors impacting development.

## 4. Methods

### 4.1. Dataset

This study includes structural magnetic resonance imaging (sMRI) data sample collected at the National Institute of Mental Health (NIMH; Bethesda, MD, USA). Participants were recruited based on their mental and physical health history for a study of brain development conducted at the NIMH between the years 1990 and 2014 [1,19,21,78]. Exclusions were based on the history of head trauma, neurological disease, and the diagnosis of psychiatric disorders. The NIMH cohort includes 2836 sMRIs acquired from participants aged 3 to 30 years, with a similar proportion of males and females across the entire age range. All participants have 1 to 7 brain scans, acquired at intervals of approximately two and a half years. This dataset is well-characterized and has been extensively described in previous works [1,19–21,78,80].

#### 4.2.1. Sample

We included two subsets of the NIMH dataset, consisting of a total of 776 participants and 1142 scans:

1. A cross-sectional sample, consisting of a total of 776 participants’ scans at their baseline age. sMRIs brain scans were acquired from typically developing individuals (357 F, 46%) from age 5 to 25 (mean age:12.4, SD:3.49) for cross-sectional analysis.
2. A longitudinal sample, consisting of 549 scans acquired from 183 participants, each being scanned at three timepoints with approximately 2.8 yrs intervals. This sample is specifically ideal for capturing maturational changes in this period (see section 4.1 for participants’ characteristics).

Other demographic and cognitive variables include IQ and SES. Details are provided in the Supplementary Materials Section S1.

### 4.2. Neuroimaging

#### 4.2.1. Image acquisition

We used T1-weighted sMRI images collected on the same 1.5 T General Electric SIGNA scanner (Milwaukee, WI), with contiguous 1.5 mm axial slices and 2.0 mm coronal slices using a three-dimensional (3D) spoiled gradient recalled-echo sequence. Image acquisition parameters are as follows: echo time: 5 ms, repetition time: 24 ms, flip angle: 45°; acquisition matrix: 256×192, number of excitations: 1, the field of view: 24 cm, resulting in a voxel resolution of 0.9375×0.9375.0×1.5 mm (for more details, see [1,19–21,78,80]). All scans underwent quality control for the raw data (Supplementary S2) and the pre-processed outputs (Supplementary S6).

#### 4.2.3. Image Pre-processing and Cortical Features Extraction

In order to standardize T1-weighted image quality, pre-processing was performed using the minc-bpipe-library pipeline (https://github.com/CobraLab/minc-bpipe-library; see Supplementary S3). These pre-processed images were then submitted to the CIVET pipeline (v 2.1.0) for cortical morphometry estimation (see Supplementary S5). SA and CT were estimated as in previous work [90,91] (blurred at 40 mm and 30 mm respectively [90]), LGI using the surface ratio [92] (using a 20 mm sphere), and finally, MC was estimated as in [93]. Further details are provided in Supplementary Materials sections S4 and S5.

### 4.3. Non-negative matrix factorization (NMF)

NMF is a data-driven matrix decomposition technique that models dominant patterns of covariance across a given dataset. The unique feature of this technique is the nonnegativity constraint which requires both input and output matrices to be non-negative. This feature of NMF is not only ideal for working with neuroimaging data as the elements of the input matrix are constrained to be non-negative but also provides an additive and parts-based reconstruction and representation of the data. These features further facilitate the intuitive interpretation of the factorization and allow for a more straightforward and biologically plausible interpretation of neuroimaging data compared to other methods of variance components that contain both positive and negative valued component weightings [44,94]. Further details are provided in Supplementary Materials section S7.

For cross-sectional analysis, the input matrix was created such that for each subject, each of the four cortical metrics’ vertex-wise measures was stacked across hemispheres to create a single column of vertex-wise data for a given subject-metric.

To investigate the coordinated pattern of cortical change across multiple measures for longitudinal analysis, we used slopes of age per subject-metric as the input to NMF. We included 183 participants with three scans (a total of 549 total scans). A linear mixed-effects regression model was performed with random intercept and slope for each subject. We next extracted subject-specific vertex-wise age-related coefficients, as a proxy of change over time, for each of the four morphometric features as the input for NMF (see Supplementary Materials S8.10 for more details).

### 4.4. Stability analysis

To select the optimal number of components (k) for OPNMF decomposition, stability analysis was performed. In order to balance the spatial stability of various decompositions with capturing major changes in the reconstruction accuracy, a split-half stability coefficient and the change in reconstruction errors were calculated for a range of 2-20 component decompositions [46] (see Supplementary S9).

### 4.5. Post NMF analysis

To assess associations between metric-wise NMF components’ weights and age, sex, IQ, and SES on the other hand, we performed multiple linear regression analyses to determine the statistical significance of the regression model (see Supplementary Materials S10.1).

To investigate possible relationships across identified individual variation patterns of morphometry and demographics data, we performed behavioral partial least square analysis (bPLS). bPLS is a multivariate technique used to capture major covariance patterns between the two sets of data via singular value decomposition (SVD) of a mutually constructed correlation matrix, which outputs uncorrelated sets of latent variables (LV). Each LV describes the linear combinations of the two datasets that are maximally covaried and can be interpreted as a pattern of association between the two sets of data [46,62–65]. (for details, see supplementary section S10.2)

### 4.6. Situating morphometric components along the gradients of brain function

We assessed how the identified morphometric covariance patterns situate along the stages of the cortical gradients’ gradual maturation (childhood, adolescence, and adulthood) by extracting the gradient value distributions of vertices within each component’s spatial boundaries from the maps of the second gradient of functional connectivity for children (6-12 years old), and the map of the first gradient map from adolescents (12-18 years old) and adults (22-35 years old). The child and adolescent maps were obtained from Dong et al. [57], and the adult map was obtained from Margulies et al. [58]. All gradient maps were transformed into CIVET space (MNI ICBM152 surface) using neuromaps toolbox [95] (https://github.com/rmarkello/neuromaps) to further assess NMF components’ position along the maps.

## Abbreviations

bPLS: behavioral Partial Least Squares
C: Component
CT: Cortical Thickness
CV: Cortical Volume
GI: Local Gyrification Index
GMV: Gray Matter volume
ICA: Independent Component Analysis
IQ: Intelligence Quotient
LC: Longitudinal Component
LV: Latent Variable
MC: Cortical Mean Curvature
NIMH: National Institute of Mental Health
NMF: Non-negative Matrix Factorization
OPNMF: Orthogonal Projective Non-negative Matrix Factorization
PCA: Principal Component Analysis
PLS: Partial Least Squares
SA: Cortical Surface Area
SES: Childhood Socioeconomic Status
sMRI: Structural Magnetic resonance imaging
T: Tesla

