## Supplementary for "Human Brain Development: a cross-sectional and longitudinal study integrating multiple neuromorphological features"

### 5. Supplementary

Table.S1. Cross-sectional and longitudinal data samples participants' demographic data characteristics.

| Characteristic \ Data sample | Cross-sectional sample | Longitudinal sample |
|---|---|---|
| Number of scans | 776 | 549 |
| Number of individuals | 776 | 183 |
| Age, years | | |
| Mean (SD) | 12.4 (3.49) | 11.2 (2.7) |
| Range | 5 - 25 | 5 - 24.2 |
| Sex, n | | |
| Female (%) | 357 (46%) | 77 (42) |
| Male | 419 | 106 |
| IQ | | |
| Mean (SD) | 110 (13.1) | 114 (12.4) |
| Range | 50 - 150 | 80 - 140 |
| SES | | |
| Mean (SD) | 44 (18.3) | 40 (19.7) |
| Range | 20 - 94 | 20 - 90 |
| Intervals between scans, years | | |
| Mean (SD) | NA | 2.8 (0.5) |
| Range | NA | 1- 6 |

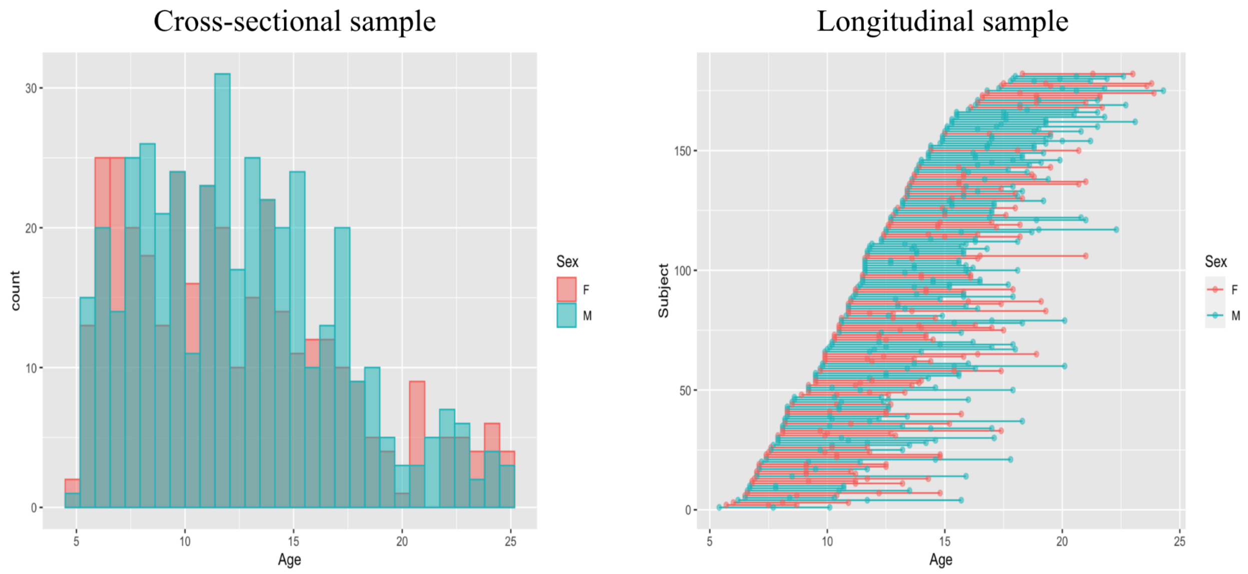


Fig.S1. NIMH subsamples characteristics. Left) Histogram showing the distribution of scans on the sample by subjects' age and sex (females in red). Right) Dot and line plot showing the distribution of scans per subject by age and sex where each line indicates a subject with repeated scans identified by dots.

**A**

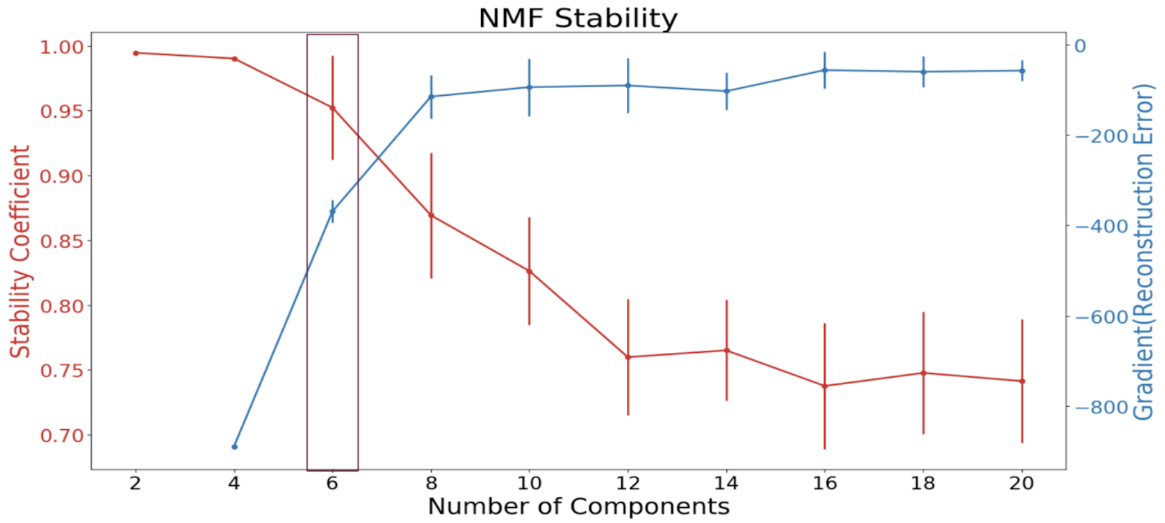


**B**

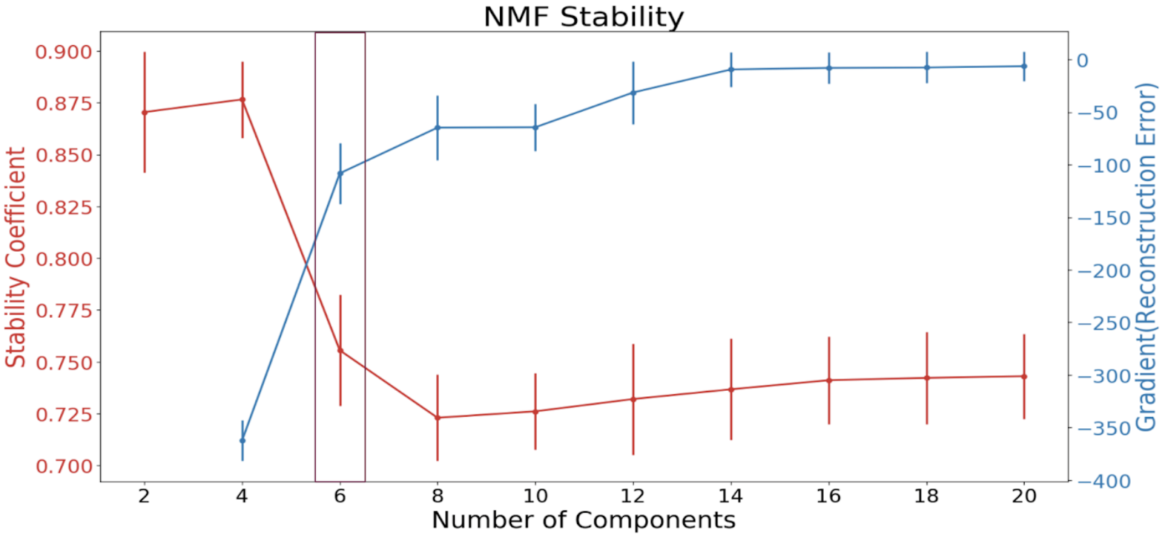


Fig.S2. A) Cross-sectional and B) longitudinal split-half stability analysis plots. The stability coefficients (red) and the gradients of the reconstruction error (blue) of OPNMF decompositions are shown for every other granularity of 2 - 20. The inverse relationship between the number of components and stability is shown. The stability (red line) drops sharply at k > 6, and the gradient in reconstruction error (blue line) from k = 6 to k = 8 is considerably less than from k = 4 to k = 6. The k = 6 decomposition solution was therefore chosen to balance the stability, and reconstruction accuracy for both decompositions.

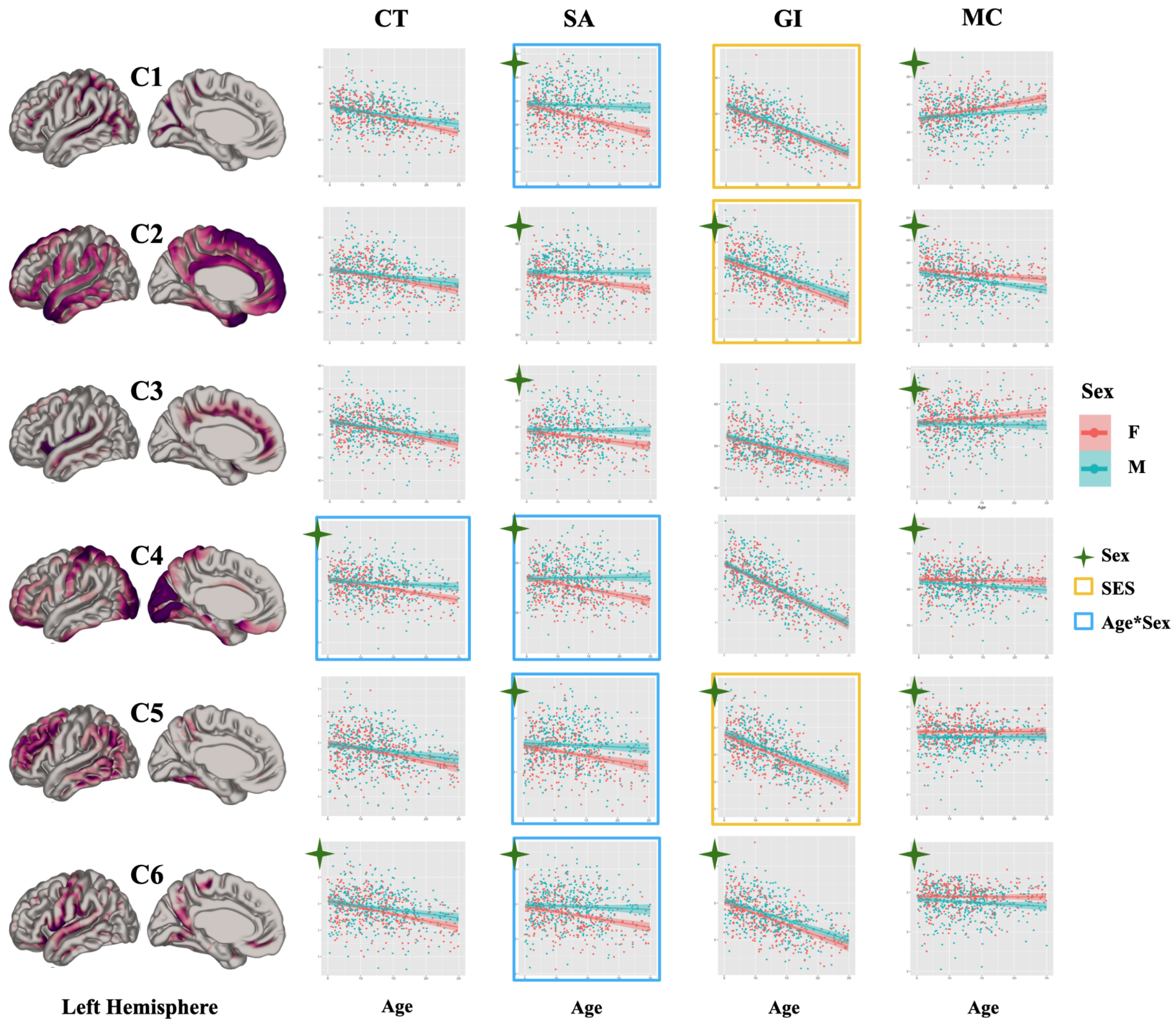


Fig.S3) Multiple linear regression models of NMF individuals' weightings corrected for multiple comparisons across components (p-value <0.008), plotted against age for males and females. Component weights with significant age-by-sex interactions are highlighted in blue, component weights significantly associated with SES are highlighted in yellow, and component weights significantly associated with sex are marked by green Astros.

Table.S2. Summary of all Cross-sectional linear model results. t-values (t) and p-values (p) (uncorrected) are bolded if they survive Bonferroni correction (p<0.008).

| Metric | CT | | | | | SA | | | | |
|---|---|---|---|---|---|---|---|---|---|---|
| factor | **Age** | **SexM** | **SES** | **IQ** | **Age*Sex** | **Age** | **SexM** | **SES** | **IQ** | **Age*Sex** |
| C1 | **t=-10.30**<br>**p=<2e-16** | t= 2.086<br>p=0.037 | t= 0.659<br>p=0.5098 | t=0.161<br>p=0.8718 | t= 2.515<br>p=0.012 | **t=-5.422**<br>**p=7.89e-08** | **t= 7.197**<br>**p=1.46e-12** | t=-1.252<br>p=0.211 | t= 0.614<br>p= 0.539 | **t= 4.125**<br>**p=4.11e-05** |
| C2 | **t= -6.842**<br>**p=1.59e-11** | t= 2.238<br>p=0.025 | t= 1.368<br>p=0.1717 | t=0.433<br>p=0.6651 | t= 1.359<br>p= 0.175 | **t= -2.854**<br>**p=0.0044** | **t= 5.777**<br>**p=1.1e-08** | t=-0.823<br>p=0.410 | t= 1.382<br>p= 0.167 | t= 2.630<br>p= 0.00870 |
| C3 | **t= -8.769**<br>**p=<2e-16** | t= 2.080<br>p=0.037 | t=0.0686<br>p=0.644 | t=0.0842<br>p=0.141 | t=1.406<br>p=0.160 | **t= -3.005**<br>**p=0.0027** | **t=5.000**<br>**p=7.11e-07** | t= -1.118<br>p=0.263 | t= 1.478<br>p=0.1397 | t= 2.453<br>p= 0.01439 |
| C4 | **t= -6.101**<br>**p=1.67e-09** | **t= 3.655**<br>**p=0.0002** | t= 1.223<br>p=0.2217 | t=0.293<br>p=0.7699 | **t=3.307**<br>**p=0.000** | **t= -3.833**<br>**p=0.0001** | **t= 6.489**<br>**p=1.55e-10** | t= 0.051<br>p=0.959 | t= 0.240<br>p=0.8107 | **t= 4.432**<br>**p=1.07e-05** |
| C5 | **t= -7.188**<br>**p=1.56e-12** | t= 1.591<br>p=0.112 | **t= 1.392**<br>**p= 0.164** | t=0.617<br>p=0.538 | t= 1.762<br>p=0.078 | **t= -4.122**<br>**p=4.16e-05** | **t= 5.674**<br>**p=1.98e-08** | t=-1.076<br>p=0.282 | t= 1.796<br>p=0.0729 | **t= 2.649**<br>**p= 0.00824** |
| C6 | **t= -8.545**<br>**p=2e-16** | **t= 2.664**<br>**p=0.007** | t= 0.924<br>p=0.3555 | t=0.249<br>p=0.8036 | **t=2.046**<br>**p=0.041** | **t=-4.428**<br>**p=1.09e-05** | **t= 6.159**<br>**p=1.18e-09** | t=-1.205<br>p=0.228 | t= 1.865<br>p=0.0626 | **t= 3.034**<br>**p= 0.0025** |

| Metric | GI | | | | | MC | | | | |
|---|---|---|---|---|---|---|---|---|---|---|
| factor | **Age** | **SexM** | **SES** | **IQ** | **Age*Sex** | **Age** | **SexM** | **SES** | **IQ** | **Age*Sex** |
| C1 | **t=-22.186**<br>**p=< 2e-16** | t= 2.297<br>p=0.0218 | **t= -2.898**<br>**p=0.003** | t=-0.575<br>p=0.5653 | t= 0.176<br>p=0.860 | **t= 6.107**<br>**p=1.61e-09** | **t= -3.170**<br>**p=0.001** | t=0.725<br>p=0.46 | t= -0.264<br>p=0.7918 | t= -2.424<br>p= 0.0156 |
| C2 | **t=-16.589**<br>**p=<2e-16** | t= 3.861<br>p=0.0001 | **t= -3.065**<br>**p=0.002** | t=0.597<br>p=0.5504 | t=0.621<br>p=0.534 | **t=-5.880**<br>**p=6.11e-09** | **t= -5.141**<br>**p=3.48e-07** | t=-0.36<br>p=0.71 | t= -0.982<br>p=0.326 | t= -1.629<br>p= 0.10379 |
| C3 | **t=-12.323**<br>**p=<2e-16** | t= 2.533<br>p=0.0115 | t= -2.543<br>p=0.0112 | t=0.360<br>p=0.7187 | t=0.877<br>p=0.380 | t= 1.548<br>p=0.122 | **t=-4.342**<br>**p=1.6e-05** | t=0.957<br>p=0.33 | t= -0.181<br>p=0.856 | t=-2.141<br>p=0.03263 |
| C4 | **t=-24.249**<br>**p=<2e-16** | t= 1.652<br>p=0.0988 | t= -2.632<br>p=0.0086 | t= -0.320<br>p=0.7487 | t=0.293<br>p=0.769 | t= -2.537<br>p=0.0114 | **t=-4.629**<br>**p=4.3e-06** | t= 0.02<br>p=0.32 | t= -0.979<br>p=0.1476 | t=-1.394<br>p=0.164 |
| C5 | **t=-19.030**<br>**p=< 2e-16** | t= 3.358<br>p=0.0008 | **t= -2.978**<br>**p= 0.002** | t= 0.011<br>p=0.9913 | t=0.306<br>p=0.759 | t= 0.056<br>p=0.955 | **t=-4.951**<br>**p=9.07e-07** | t=-0.10<br>p=0.91 | t= -1.377<br>p=0.169 | t= -0.122<br>p=0.9032 |
| C6 | **t= -17.391**<br>**p=< 2e-16** | t= 3.404<br>p=0.0006 | t= -2.104<br>p=0.0357 | t= 0.312<br>p=0.7554 | t=0.981<br>p=0.327 | t= -1.756<br>p=0.0795 | **t=-5.101**<br>**p=4.25e-07** | t=0.074<br>p=0.94 | t= -0.208<br>p=0.8352 | t= -1.515<br>p= 0.130 |

Table.S3. Summary of all longitudinal NMF linear model results. t-values (t) and p-values (p) (uncorrected) are bolded if they survive Bonferroni correction (p<0.008).

| Metric | **CT** | | | | | **SA** | | | | |
|---|---|---|---|---|---|---|---|---|---|---|
| factor | **Age** | **SexM** | **SES** | **IQ** | **Age*Sex** | **Age** | **SexM** | **SES** | **IQ** | **Age*Sex** |
| C1 | t=--0.527<br>p=0.5991 | t= 2.086<br>p=0.0183 | t= 0.807<br>p= 0.4210 | t= -0.274<br>p= 0.7847 | t= 1.740<br>p=0.0836 | t=-2.371<br>p=0.0188 | t= -0.028<br>p=0.9775 | t=1.469<br>p=0.1437 | t= -2.154<br>p= 0.0326 | t= 2.638<br>p=0.009083 |
| C2 | t= -2.051<br>p=0.041715 | t= 3.610<br>p=0.0145 | t= 0.580<br>p=0.56266 | t=0.613<br>p=0.54047 | **t= 2.677**<br>**p= 0.008** | t= -1.237<br>p=0.2176 | t= 0.130<br>p=0.8963 | t=-1.252<br>p=0.2124 | t= -2.159<br>p= 0.0322 | **t= 3.037**<br>**p= 0.00275** |
| C3 | t= 0.245<br>p=0.8069 | t= 2.469<br>p=0.037 | t=0.687<br>p=0.4927 | t=0.0842<br>p=0.141 | t=1.707<br>p=0.0896 | t= -2.605<br>p=0.00996 | **t=2.784**<br>**p=0.00596** | t= -1.118<br>p=0.1521 | t= 0.780<br>p= 0.4366 | t= 2.458<br>p= 0.014935 |
| C4 | t= 0.067<br>p=0.94644 | t= 2.641<br>p=0.00899 | t= 0.496<br>p=0.62020 | t=0.268<br>p=0.78899 | t=1.270<br>p=0.206 | **t= -3.654**<br>**p=0.000340** | **t= 4.904**<br>**p=2.11e-06** | **t= 3.370**<br>**p=0.0009** | t= -0.132<br>p=0.89522 | t= 2.518<br>p=0.012691 |
| C5 | t= -0.049<br>p= 0.9609 | t= 1.813<br>p=0.0715 | t= 1.392<br>p= 0.164 | t=1.159<br>p=0.2479 | t= 0.682<br>p=0.496 | t= -0.581<br>p=0.5618 | t= 0.539<br>p=0.5904 | t=-1.369<br>p=0.1728 | t= 1.949<br>p=0.0529 | t= 1.120<br>p= 0.2644 |
| C6 | t=0.486<br>p=0.6279 | t= 1.862<br>p=0.0643 | t= 0.421<br>p=0.6746 | t=0.151<br>p=0.8799 | t=1.790<br>p=0.0752 | t=--2.182<br>p=0.0304 | t=1.249<br>p=0.2132 | t=-1.815<br>p=0.0713 | t= -0.765<br>p=0.4451 | t= 1.018<br>p= 0.3103 |

| Metric | **GI** | | | | | **MC** | | | | |
|---|---|---|---|---|---|---|---|---|---|---|
| factor | **Age** | **SexM** | **SES** | **IQ** | **Age*Sex** | **Age** | **SexM** | **SES** | **IQ** | **Age*Sex** |
| C1 | t=1.413<br>p=0.1596 | t=-1.085<br>p=0.2794 | t= 0.807<br>p=0.4210 | t=-2.076<br>p= 0.0394 | t= 0.598<br>p=0.5504 | t= 2.083<br>p= 0.0387 | t= 2.404<br>p= 0.0173 | t=-2.001<br>p= 0.046 | t= 1.695<br>p=0.0919 | t= -2.285<br>p= 0.02353 |
| C2 | **t=4.094**<br>**p=6.43e-05** | t= -0.608<br>p=0.5442 | t= 1.028<br>p=0.3052 | t=--1.802<br>p= 0.0732 | t=-0.388<br>p=0.6985 | t=0.415<br>p=0.67863 | **t= 3.138**<br>**p=0.00199** | t=-1.633<br>p=0.104 | t= 1.559<br>p=0.12075 | **t= -3.239**<br>**p= 0.00143** |
| C3 | t=1.823<br>p=0.0700 | t= -0.328<br>p=0.7433 | t= 0.229<br>p=0.8190 | t= -1.763<br>p=0.0796 | t=-0.071<br>p=0.9431 | t=0.091<br>p=0.9275 | t=2.198<br>p=0.0293 | t=-1.086<br>p=0.278 | t= 0.254<br>p=0.7995 | t=-0.990<br>p=0.324 |
| C4 | **t=4.571**<br>**p=9.08e-06** | t= -0.828<br>p=0.409 | t= 0.794<br>p=0.428 | t= -1.343<br>p=0.181 | t=0.190<br>p=0.8498 | t= 0.906<br>p= 0.366 | t=0.847<br>p= 0.398 | t= -0.86<br>p= 0.390 | t= 1.071<br>p=0.286 | t=-0.871<br>p=0.385 |
| C5 | t=-0.784<br>p=0.4344 | t= -0.018<br>p=0.9856 | t= -0.640<br>p= 0.5231 | t=-1.859<br>p=0.0646 | t=0.116<br>p=0.9076 | t=-0.598<br>p=0.550 | t=0.474<br>p=0.636 | t=-0.854<br>p=0.394 | t=0.091<br>p=0.928 | t= -1.372<br>p=0.172 |
| C6 | t=0.030<br>p=0.976 | t= 0.565<br>p=0.573 | t= 1.319<br>p=0.189 | t= -1.620<br>p=0.107 | t=0.661<br>p=0.509 | t= 2.331<br>p=0.0209 | t=-0.244<br>p=0.8072 | t=-1.796<br>p=0.074 | t= 0.646<br>p=0.5189 | t= -0.706<br>p= 0.4813 |

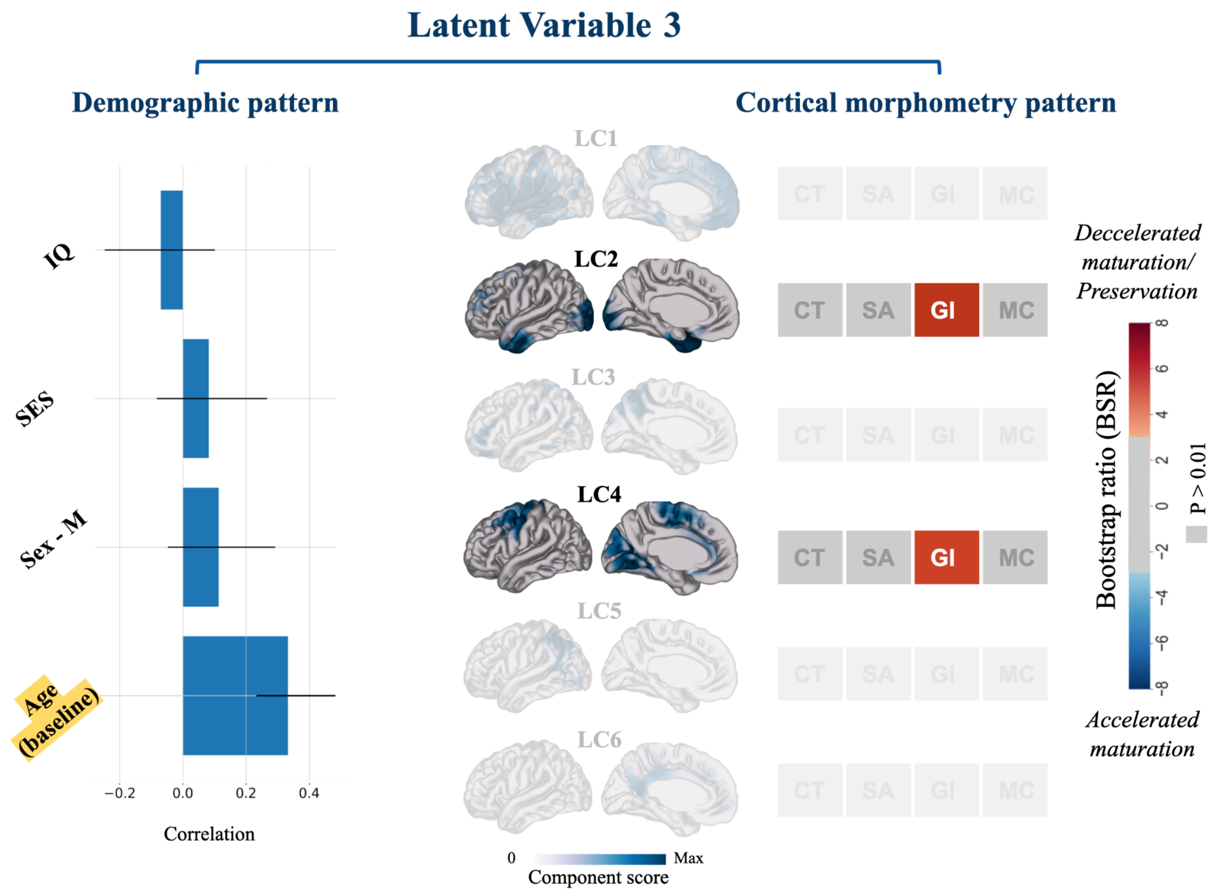


Fig.S4) LV3 (p = 0.02), explaining 24% of covariance across the data (p-value = 0.005) describes a specifically age-related pattern in which changes of GI in two cortical components of LC2 and LC4, such that GI slopes show a milder decline in older ages (the slopes of GI change become less negative in older ages) (Fig.S4). Together, these LVs (explaining 97% of the covariance) uncover patterns of co-ordinated anatomical change that are sexually differentiated and influenced by environmental factors such as SES and related to differing cognitive abilities.

### S1. Demographic and cognitive variables

1) Full-scale Intelligence Quotient [IQ] estimation, using an age-appropriate Wechsler scale. 94% of participants have been assessed with the Wechsler Abbreviated Scale of Intelligence (WASI) [96]; the rest include WAIS-R, WISC-R, WISC-III, WPPSI, and WPPSI-III. Notably, since most individuals with repeated scans had fewer IQ estimated measures, we used each participant's most recent IQ estimation for the longitudinal sample analysis [78].

2) Childhood socioeconomic status (SES) score quantified by the Amherst modification of the Hollingshead two-factor index [97,98]. This assessment is based on parental education and occupation, which was used to obtain a single SES score for analysis. Notably, the conventional directionality suggests that a lower Hollingshead score indicates higher SES. The lowest Hollingshead score (i.e., 20) corresponds to individuals from the most advantaged families, while those from the least advantaged families have the highest Hollingshead score (i.e., 140) [78]. Accordingly, the Hollingshead score that has been used in the statistical analyses and figures has an opposite direction with socioeconomic status. For ease of interpretation, the directionality of socioeconomic status itself (rather than the Hollingshead score) will be discussed in the rest of the present work. Demographic and cognitive characteristics of the NIMH dataset have been extensively described in [78].

#### S2. Raw scan motion quality control

MRI data are prone to be affected by in-scanner head motion, involuntary movements, or physiological sources of noise such as cardiac cycle and respiration, which subsequently may degrade the image quality and lead to general and regionally-specific biases and misinterpretation of the quantitative outputs derived from these images. Due to the inverse relationship reported between in-scanner motion and age of the participants [99], the issue is more critical in studies involving younger aged populations and children, such as the present work. In the present work, all raw images were visually assessed and quality controlled for motion artifacts, such as ghosting and blurring, using the QC procedure previously developed in our group and described in [59,100] (for details, see: https://github.com/CoBrALab/documentation/wiki/Motion-Quality-Control-Manual).

#### S3. minc-bpipe-library pipeline

This pipeline, by default, performs the integrated steps of N4 bias field correction to correct for intensity inhomogeneities [101], image registration to Montreal Neurological Institute (MNI) space (ICBM 2009c Nonlinear Symmetric) using bestlinreg [102,103], cropping the neck, field-of-view standardization using an inverse-affine transformation of an MNI space head mask, brain extraction and generating a brain mask using Brain Extraction based on nonlocal

Segmentation Technique (BEaST) [104] as previously described in [105]. Following the preprocessing steps, the extracted brain bias field corrected T1 in native space with a specific brain mask for each subject was submitted to the CIVET processing pipeline.

#### S4. Surface-based morphometry feature estimation

To estimate quantitative neuroanatomical features, we used the CIVET pipeline (Version 2.1.0; Montreal Neurological Institute; http://www.bic.mni.mcgill.ca/ServicesSoftware/CIVET; [90,106,107]. Preprocessed T1 Weighted sMRI scans with matched masks, generated in minc-bpipe-library, were submitted to the CIVET for automated surface-based estimation of four cortical features of CT, SA, GI, and MC at 81,924 vertices across the cortex. T1w images were linearly registered to the Montreal Neurological Institute (MNI) ICBM 152 average [108], brain tissue was classified into white matter, gray matter, and cerebrospinal fluid [109], and the surface was extracted using the Constrained Laplacian Anatomical Segmentation using Proximities (CLASP) method [90,91].

#### S5. Extracting morphometric features

Followed by surface extraction, neuroanatomical metrics were estimated as follows:

**CT:** Cortical thickness was defined and estimated as the minimum distance between the gray matter and white matter surfaces at each vertex [61,110]. CT maps were blurred using a 30-mm full-width at half-maximum (FWHM) surface-based diffusion smoothing kernel [90].

**SA:** The surface area was calculated by the Voronoi method, in which the subject area is divided by the model area at each vertex on the surface template on an intermediate tessellated surface mesh between pial and gray/white surfaces [111]. The SA maps were blurred using a 40-mm geodesic surface kernel [90].

**GI:** Local Gyrification Index was calculated for the local estimation of cortical folding at each vertex as the ratio between the pial surface contained in a sphere placed at the vertex and the surface area of a circle of equivalent radius [92,112]. GI measures were estimated at a 20 mm radius.

**MC:** Mean cortical curvature was calculated as the average of principal curvatures, derived from the inverse of the radius of the osculating circles at each vertex on the mid surface of the gray and white matter junction [93].

Notably, the non-cortical midlines were masked out for analyses (4802 vertices), and we proceeded with the morphometric data of 77,122 vertices for each subject.

#### S6. Output quality control

CIVET automatically produces figures showing the gray and white matter classification and boundary delineations for quality control. To prevent misleading quantitative results in further analyses [113], all CIVET outputs were controlled for white and gray matter classification accuracy and surface delineation by visual inspection following the QC procedure previously developed in our group [59]. Only scans that passed through the CIVET quality control will be used to extract morphometric features to be included in our analyses.

#### S7. Non-negative Matrix Factorization

In the current work, we employed the orthonormal projective variant of NMF (OPNMF) [44,114], which prioritizes sparsity in the solution and the corresponding part-based decomposition. The goal is to provide minimally overlapping components where orthogonality ensures each vertex is easily assigned to a particular component for a purely additive parts-based representation and improves the specificity, and the projective features ensure that all components participate in the reconstruction of the input data sample, which improves sparsity.

Orthogonal Projective NMF background:

OPNMF decomposes an input matrix V of dimensions [m x n] into a component matrix W (m x k) and a weight matrix H (k x n) in which k is the number of components pre-defined by the user (see Methods section 3.6). Decomposition of the input matrix is such that the multiplication of the component and weight matrices reconstructs the original input as accurately as possible with the minimum reconstruction error between the original input and the reconstructed input (W x H) (Fig.S5) [44,46,115]. The W and H matrices can be used to describe patterns of variance across both axes of the input matrix.

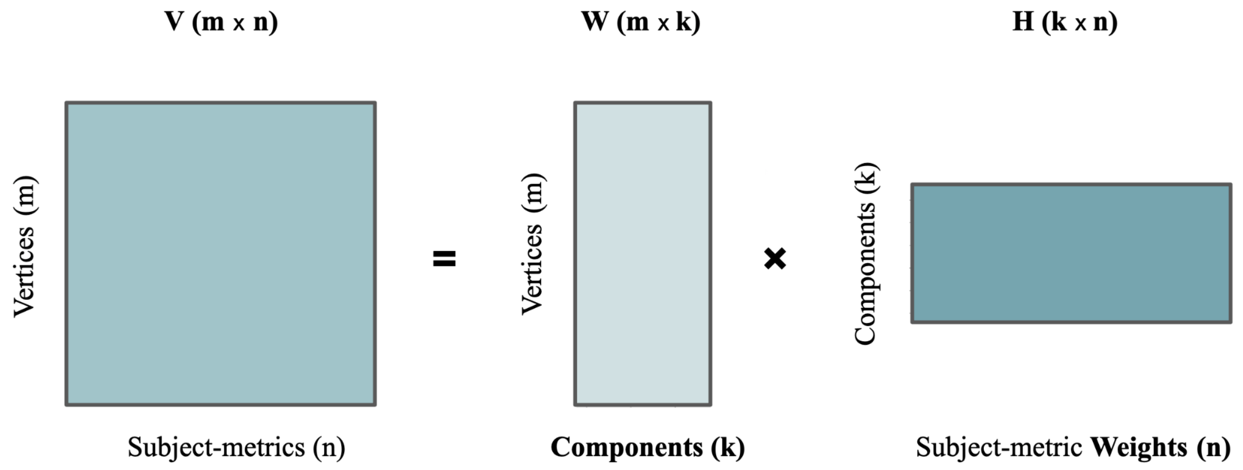


Fig.S5. A representation of non-negative matrix factorization (NMF). NMF decomposes an input matrix such that the multiplication of the two outputs reconstructs the input matrix with the minimum reconstruction error. (adapted from Patel et al. [46])

#### Cross-sectional implementation of NMF

Given the inclusion of 776 subjects and the columnar stacking fashion of the four morphometric features of CT, SA, GI, and MC, the input matrix was built having 776* 4 columns such that the first 776 rows correspond to subjects' CT values, second 776 rows correspond to subjects' SA values, etc. Finally, a within-subject z-scoring normalization was performed to have different metrics with varying magnitudes on the same scale and shifted by the minimum z-scored value to eliminate negative values. This multivariate morphometry matrix would then be submitted to OPNMF.

#### Running OPNMF

Following the construction of the input matrix, the OPNMF run was performed using the publicly available code by [44,114,116,117] at (https://github.com/asotiras/brainparts) using Octave (https://www.gnu.org/software/octave/doc/v5.2.0/). The algorithm was initialized with a non-negative, double singular value decomposition (SVD) and followed a maximum iteration of 100000 and tolerance = 0.00001, as previously described in [46].

#### Interpreting OPNMF outputs

OPNMF outputs a components matrix (W) and weights matrix (H), which together reconstruct the input matrix:

1) Component matrix (W) of dimensions [cortical vertices (m) x number of components (k)]: indicates the spatial location of the components indicating the extent to which each vertex in the brain loads onto each component. Due to the orthogonality feature of OPNMF, clustering can easily be derived from OPNMF using a winner take all approach, where each vertex is assigned to the cluster number corresponding to the component in which it has the highest weight. Mapping back these identified clusters of vertices to the population average brain enables visualization of a parts-based representation of spatial cortical components that share the same patterns of covariance across four morphometric features.

2) Weight matrix H of dimensions [number of components (k) x subject-metric pairs (n)] represents each subject-metric pair contribution and loading onto each of the identified components. Accordingly, a higher subject-metric loading onto a given component's vertex would indicate a greater magnitude of that metric within the spatial area specified by the component. The components and weight matrix are then jointly used to describe the spatial location of identified components (W matrix) and the pattern of covariance across multiple morphometric features of the cortex (H matrix).

#### S8. Longitudinal implementation of NMF

To extract subject-specific cortical slopes for each metric, a vertex-wise linear mixed-effects model was performed with age as the fixed effect and a random intercept and slope of age for each subject. Models were implemented in R (Version 3.6.3; www.r-project.com).

The coefficients for *j*th subject's *i*th vertex's metric *n* from time-points 1 to 3 were calculated based on the regression model as follows:

$$Y_{subjectj\_vertexi\_metricn} = \beta 0 + dij + \beta 1\ (Age) + \beta j\ (1+Age|Subject) + eij \quad (1)$$

Where Y corresponds to subjects' metric measures at each vertex, $\beta_0$ corresponds to equation intercept, $\beta_1$ corresponds to fixed-effect coefficient, and $\beta_j$ corresponds to subjects' random B-value. From each model, subject-specific age-related slopes (i.e. coefficient) were extracted using the R coef() function. Coefficients are the summations of the general fixed effect of age ($\beta_1$) and subject-specific random effects ($\beta_j$) [118], representing the impact of age on each cortical measure. Finally, to capture patterns of 'coordinated change' across cortical features, the input matrix was reconstructed similar to cross-sectional methods (see section 3.4.1) with columns containing vertex-wise calculated coefficients of age in three consecutive timepoints for each subject-metric pair (z-scored and shifted by the minimum value).

#### Interpreting longitudinal OPNMF outputs

Similar to outputs described in section 3.4.4, longitudinal OPNMF outputs a component and a weights matrix that jointly describe different covariance patterns of longitudinal change. In the weight matrix, higher weights indicate a relatively higher magnitude of slopes, implying a slower decline, preservation, or decrease in the specific metric. Lower weights indicate a lower magnitude of slopes, implying relatively sharper decline and loss in the specific morphometric feature.

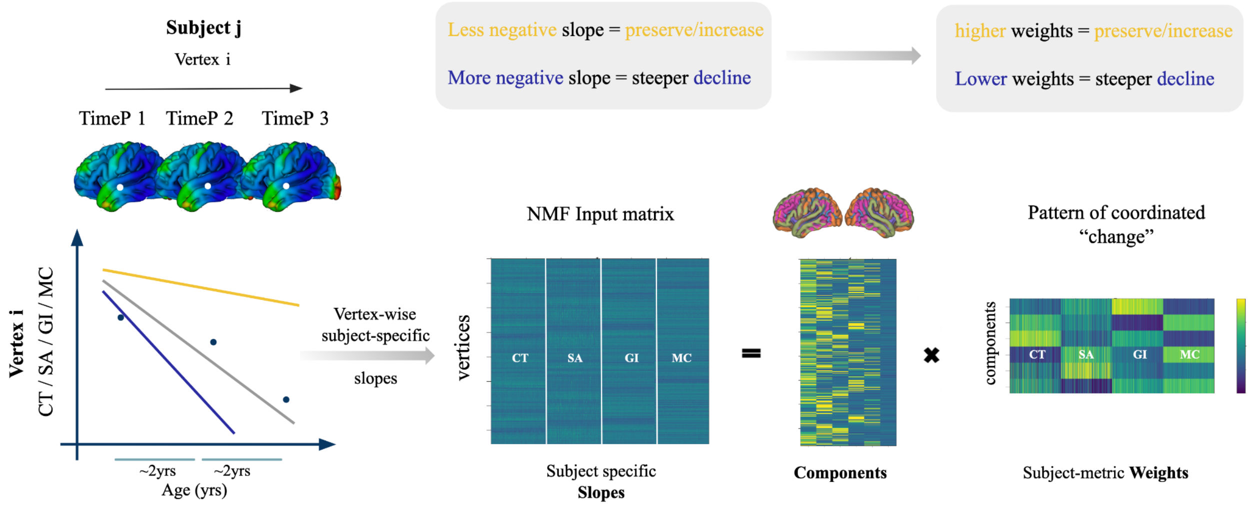


Fig.S6. A schematic representation of longitudinal NMF implication and interpretation. While higher NMF weights indicate relative preservation of a metric, a lower weight indicates a steeper decline over time.

#### S9. Stability Analysis

To assess the stability, we measured the spatial similarity between the component's output of OPNMF runs across varying splits of the input data at each granularity of 2 - 20. Accuracy was calculated by reconstruction error, and we plotted the gradient or change in the reconstruction error from one granularity to the next (k to k + 1) [46] to identify granularities that capture the largest patterns of variance. The procedure was done by splitting the participants (n=776) into two groups of a (n_a = 388) and b (n_b = 388) ten times. A separate four-metric input matrix was built (Va and Vb) for each half-split, and we performed OPNMF on each of the half-splits independently. Each run outputs a weight and a component matrix, Wa and Wb matrices, of dimensions [#vertices * k].

To calculate the stability, a similarity matrix for each of Wa and Wb matrices (c_Wa, and c_Wb) was computed using cosine similarity for rows of W matrices such that in each similarity matrix, the cosine similarity of component scores between a specific vertex and all other vertices is represented in each row. The higher the cosine similarity for each vertex with other vertices, the more similar the component scores between a pair of vertices. Next, the correlation coefficient between corresponding rows of the similarity matrices of each split (c_Wa, and c_Wb) was computed. The mean correlation across all rows (vertices) was taken as an indicator for stability at that specifically tested granularity such that a mean correlation coefficient of 1 represents the highest stability, while −1 represents the lowest stability [46]. These procedures were repeated for ten random splits of the data and at each even granularity of k = 2 - 20 [46].

#### S10. Post NMF analysis

##### S10.1. Multiple linear regression modeling

To this end, we ran a metric paired components-wise regression model, in which for *i*th components' *j*th metric was modeled as follows:

$$Y_{Component\ (i)\_metric\ (j)} = \beta_0 + \beta_1(Age) + \beta_2(Sex) + \beta_3(IQ) + \beta_4(SES) \qquad (2)$$

$$Y_{Component\ (i)\_metric\ (j)} = \beta_0 + \beta_1(Age \times Sex) + \beta_2(IQ) + \beta_3(SES) \qquad (3)$$

Where Y corresponds to metric-wise OPNMF components weights, $\beta_0$ corresponds to equation intercept, and $\beta_i$ corresponds to fixed-effect coefficient. We looked at all regression beta coefficients, with particular emphasis on age effects grouped by sex, age, and age and sex interactions. We looked for associations between component weights and each demographic variable, and corrected for multiple comparisons across components with Bonferroni correction [119] thresholded at a significance of p-value < 0.008

#### S10.2. Partial least squares analysis

For each LV, a singular value is computed describing the percentage of the data explained by that LV. Within each LV, both sets of variable parameters will be attributed to scores, reflecting the extent to which each parameter contributes to the identified covariance patterns represented by LVs [46,62–65]. In the context of neuroimaging analysis, this method has been originally and conventionally used to relate a set of neuroimaging data (i.e., voxel- or vertex-wise data) to a set of behavioral data [62–64]. This method has been further developed in our previous works to enable relating integrative structural and morphometric patterns derived from NMF, to inter-individual differences [46,55]. Here, we performed behavioral PLS to assess patterns of correlation across two sets of 1) brain data, the H matrix output from OPNMF containing components' subject-metric pair weights, and 2) demographics and cognitive data, including subjects' age, biological sex, SES, and IQ scores. The brain matrix of dimensions (776 subjects × 4 metrics × 6 components) and a demographic matrix of dimensions (776 subjects × 4 demographic variables) were used as the input matrices of PLS (Sex coded as 0 = Female, 1= Male, Age measured in years, SES measured in Hollingshead two-factor index, and IQ measured with Wechsler Intelligence Scales). PLS analysis outputs LVs representing patterns of covariance across component-wise morphometric brain data and demographic data. To assess the statistical significance of output LVs, we performed 10000 permutations testing such that rows of the brain data matrix were permuted 10000 times to obtain a null distribution of singular values under the assumption of permutations eliminating existing brain-demographics correlations, which yields the computation of a non-parametric P-value for each LV in the primary data (non-permuted) [46]. We then applied a $P<0.05$ threshold for consideration of LVs significance, indicating a chance of a 95% confidence that the singular value of the primary LV exceeds that of a singular value of the permuted LV [46]. To assess the contribution of each brain variable (component-wise morphometric brain data) to each identified LV, we employed bootstrap resampling. We randomly sampled 10000 sets of each brain and demographics matrices and replaced the rows to create a distribution of the singular vector weight of each brain variable in each LV. Next, to examine the contribution and reliability of a given brain variable, the bootstrap ratio (BSR) was calculated as the ratio of generated singular vector weight over the standard error of the weight from the bootstrap distribution. We then applied a threshold of $P<0.01$ (99% confidence) for consideration of brain variables' contribution significance, corresponding to a $BSR<2.58$ [46,62–65].
